# Phase transitions in a simple model of focal stroke imitate recovery and suggest neurorehabilitation strategies

**DOI:** 10.1101/2022.12.14.520421

**Authors:** Alba Carballo-Castro, Luís F Seoane

**Affiliations:** Facultad de Matemáticas, Universidad de Sevilla, C/ Tarfia s/n, 41012 Sevilla, España; Departamento de Biología de Sistemas, Centro Nacional de Biotecnología (CSIC), C/ Darwin 3, 28049 Madrid, Spain; Grupo Interdisciplinar de Sistemas Complejos (GISC), Madrid, Spain

**Keywords:** stroke, brain plasticity, brain reorganization, bilateral symmetry, neurorehabilitation

## Abstract

A stroke is a brain insult that can take offline (often permanently) extended regions of the brain. As a consequence, cognitive tasks or representations implemented by the affected circuitry lose their computational substrate (they become *orphan*). The brain must adapt to attempt retaining such functions. The existing clinical literature offers a complex picture, often with conflicting observations, about how the brain gets reorganized after stroke. It also does little use of the few mathematical works on the topic. Can a minimal mathematical model of cortical plasticity shed light on this complex phenomenology? Here we explore such minimal model, and find a specific phenomenology: a lasting perilesional reorganization for small injuries, and a temporary contralesional reorganization for large injuries that is not always reverted to ipsilesional. We furthermore show the mechanisms behind these dynamics in our model: a second order phase transition with a critical point, as well as a delayed engagement of perilesional reorganization in large injuries. These dynamics emerge out of a fairly minimal modeling of plasticity, and they reproduce the story put together from clinical observations. We further explore neurorehabilitation strategies, and argue that increased tissue susceptibility (a property that diverges at critical points) can be crucial to manipulate plasticity in beneficial ways.

## I. INTRODUCTION

Strokes are a major health concern world-wide. In the European Union alone, it affected over 1 million people in 2017, resulting in above an accumulated 9 million survivors and almost half a million deaths. These trends are expected to worsen over the next decades [1]. Strokes are caused by the disruption of blood flow, which leads to a temporary or permanent loss of specific cortical circuitry. The topography (location) and extent of the lesion determine which brain areas are disrupted, as well as whether and which cognitive abilities are affected. Some common outcomes are aphasia and motor impairment. Trivially, strokes affecting larger areas tend to cause greater cognitive deficits [2–11].

Brains are highly plastic organs [12, 13], with much of their flexibility surviving into adult age [14– 18]. Plasticity allows our brains to withstand drastic rearrangements—e.g. as required following hemispherectomy [19–21] or after a stroke. Great speculation exists concerning the phenomenology of brain reorganization after a stroke [8, 9, 11, 22–27]. Available clinical studies present a complex picture that changes across patients, over time after lesion, and with injure size and topography. Some principles seem to emerge: (i) Both perilesional and lesion-contralateral, homologous regions can be recruited. (ii) Increased activity in homologues might be due to compensatory mechanisms—e.g. use of the left hand after the right one became paretic. (iii) Contralateral recruitment is observed in aphasia as well [23, 24]. This suggests not a compensatory mechanism (as there are not ‘left’ and ‘right’ languages), but an actual involvement of distant homologues in function recovery. (iv) Recruitment might result from disinhibition—i.e. a damaged lateralized circuit becomes unable to block activity in the contralateral or perilesional tissues. This is consistent with a view of neural circuits ‘competing’ to carry out function in response to suitable inputs. (v) Contralesional circuitry might also be engaged as a relay, to distribute signals over ipsilesional areas [23].

All these possibilities appear entangled in each specific patient, and their contribution to brain reorganization might vary as recovery progresses. Concerning recovery, a compelling story, summarized in Fig. 1**a-e**, emerges out of the intricate clinical evidence [8, 9, 11, 23, 24]: (i) After small injuries, perilesional circuitry takes care of any lost functionality. But following a large enough trauma, contralateral homologues of the damaged tissue take over. This would provide a relief to the injured circuits, which could focus on healing. (ii-a) If damage is not permanent, the affected areas would come back online, and control would be returned to them. (ii-b) If some circuitry has been lost for good, control can still be returned to the perilesional region—which must adapt to host the orphaned function. (iii) But, either after temporary or permanent damage, control is not always returned to ipsilesional circuits. A viewpoint holds that sustained contralateral activity is maladaptive or detrimental to functional recovery—which has been proved in some cases [11, 24, 28]. The alternative (contralateral activity being necessary for improved function) has also been observed [26, 29].

**FIG. 1.**
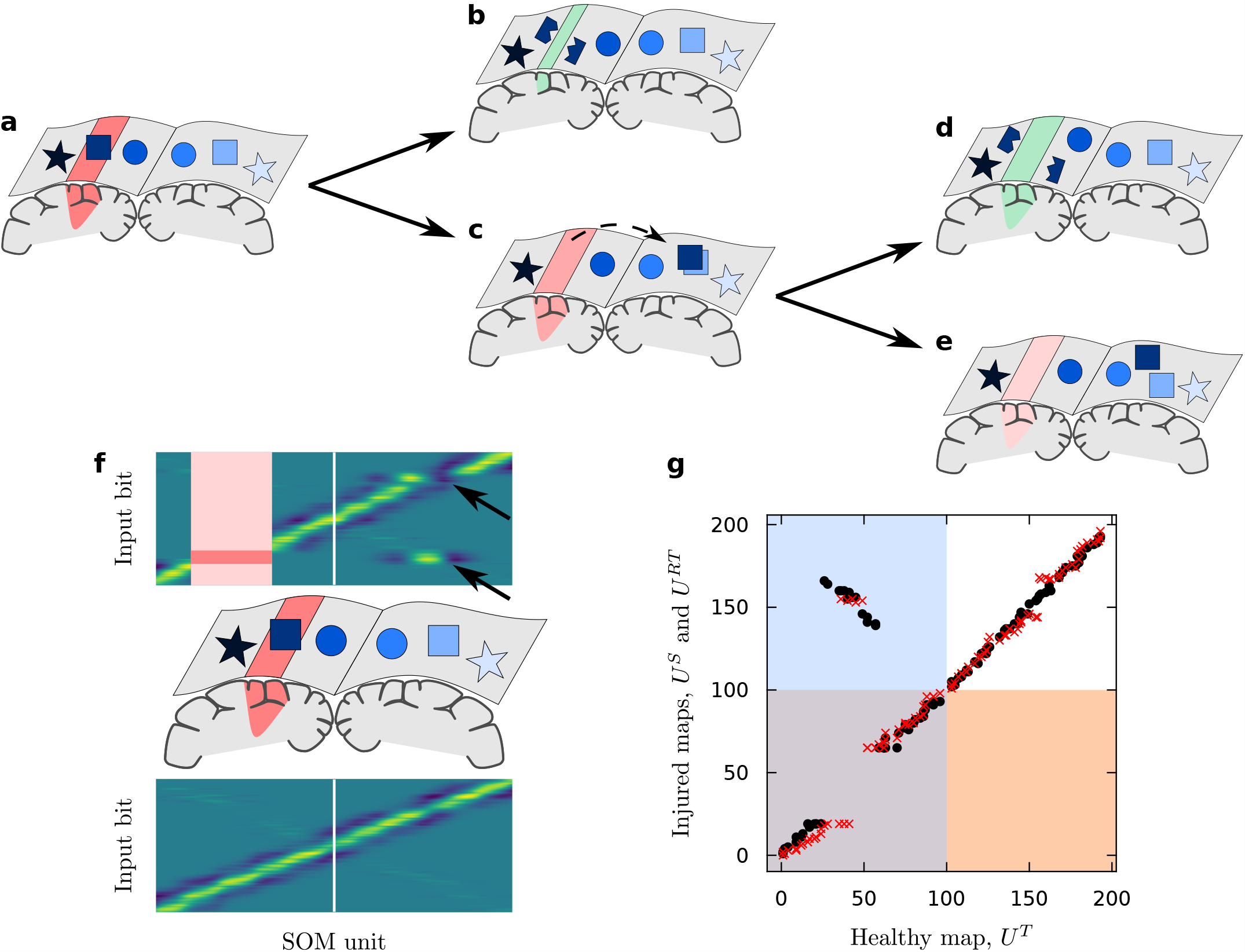
Brain reorganization after stroke. **a-e** Dynamics inferred from clinical observations and reproduced by the model. **a** A stroke takes a cortical region offline (red), resulting in an *orphan* representation (square—this would stand for, e.g., a hand becoming paretic because the area controlling it got damaged). **b** If the affected region is small enough, orphaned signals are split and remapped into perilesional circuitry. **c** If the damage is large enough, immediately after stroke the homologue-contralesional regions take care of orphan function. **d** With enough time, orphan signals might get remapped into ipsilesional areas. This is often viewed as convenient for recovery. **e** By chance, contralesional circuitry might retain control. This can be viewed as harmful or maladaptive. **f** Weights of a stroke before (bottom) and after (top) a stroke and retraining. Vertical white lines indicate brain halves. The stroked area is marked with a light red shading. Within it, a darker shading indicates orphan input stimuli. These, after retraining, appear mostly represented contralesionally (bottom arrow), which results in diaschisis (top arrow). **g** Further exemplifying reorganization in our model. We represent where each specific input was mapped by a healthy map before stroke (horizontal axis) and by a damaged map (vertical axis) immediately after stroke (black dots) or after stroke and retraining (red crosses). The blue shading represents the simulated left hemisphere (first *N/*2 neural units) in the healthy map. The orange shading represents the simulated left hemisphere in the damaged maps. Tracking whether dots or crosses fall within or outside the shadings tells us whether representation became contralateral (upper-left quadrant) or remained ipsilateral (lower-left quadrant).

Again, we are dealing with a very complex picture that results, among others, from the interplay between short- versus long-range brain plasticity, redundancy of the neural computational substrate (as provided by bilateral symmetry or local backups), and injury size. While seemingly contradictory pieces of clinical evidence accumulate, we attempt to make some progress with minimal mathematical models. What are the simplest reorganization dynamics that emerge given some barest account of brain plasticity? How does each of the relevant elements (bilateral symmetry, local versus global plasticity, insult extent) affect the reorganization outcome? Can we design interventions that accelerate neurorehabilitation?

We tackle these questions by building on a rather minimal model of brain reorganization after hemispherectomy [20]. This model uses Self-Organized Maps (SOM) [30] as an abstraction of spatially-sorted feature maps (such as the somatosensory cortex) or of competing structures across hemispheres (such as language domains, which start out as bilaterally-symmetric and become lateralized as language matures [31–33]). This minimal model ignores much of the existing complexity of these regions, focusing instead on the interplay between conflicting topological arrangements. More explicitly, feature maps are sorted as responding to stimuli along a one-dimensional segment; and a variable degree of bilateral symmetry is introduced that interferes with this linear order (Fig. 1**f**). In [20], a SOM is trained to respond to input signals with a given topology (which then the SOM acquires). An hemispherectomy is simulated by removing half of the map, leaving some input signals *orphan*. Further training leads to brain reorganization guided by either topological schema (linear-lateralized versus bilateral). This results in insights about favored plasticity pathways and window periods.

To simulate a stroke we perform the same procedure; but we damage a limited region fully contained within one hemisphere (Fig. 1**a**). We study reorganization as a function of time after stroke and of injury size, and for permanent and temporary damage. We try out neurorehabilitation strategies. As an indicator of sustained pathology, we measure the likelihood and extent of contralesional SOM representations. As noted above, in real scenarios, sustained contralesional activity is not necessarily harmful—with evidence for and against. But having protocols to prompt or prevent contralateral representations is a step towards manipulating plasticity. Having a mathematical model further allows us to ascribe causality to the elements behind our intervention, and to understand why each outcome is observed. Some previous models of neural reorganization after a stroke focus on rather local reorganization [34–39]. A notable exception are works by James Reggia and colleagues [40–44], which we discuss below and which explicitly model interhemispheric communication.

We approach this problem inspired by statistical mechanics. Our input signals and SOM were purposefully designed to resemble spins, which interactive dynamics are known to contain minimal, yet universal computational capabilities [45]. In our framework, phenomenology is often guided by phase transitions, critical points, symmetry or symmetry-breaking [20, 47–50], and other ingredients for which statistical mechanics offers a very solid bedrock. As we will show, some of these concepts constitute the key mechanisms that enable our model to reproduce clinical observations.

In Sec. II we introduce our mathematical model and sketch our simulated neurorehabilitation protocols. Secs. III.A and III.B study spontaneous reorganization after a simulated stroke that causes a permanent lesion. We observe how immediate reorganization is preferably ipsilateral for small injuries and contralateral for large ones. A sharp threshold marks this divide, reminding us of a second-order phase transition. Tracking SOM activation over time results in the trend observed in clinical studies (initial displacement of activity to contralateral homologues, gradual recovery of ipsilateral control). This pathway to recovery is not always taken—neither in the clinic, nor in our model. By chance, contralesional homologues might retain control. We understand this as a pathological condition and, in Sec. III.C, we try interventions to mitigate it. In Sec. III.D we study reorganization after temporary lesions. The tradeoff between local versus contralateral reorganization further interacts with the lesion timing. Counter-intuitively, this results in a worst-case lesion duration that reveals a delayed onset of perilesional reorganization. In Sec. IV we provide an extended discussion framing our findings in clinical observations and neurorehabilitation literature.

## II. METHODS

### A. Modeling topographic maps

Our model is a slight variation of the one in [20] to investigate hemispherectomy. Here we paraphrase its description. We refer the reader to [20], where some parameters are explored and some choices are justified. The most relevant decisions concern the topological arrangement of input signals. Inspired by the disposition of the topographic representation of our body, we are interested in inputs, and maps, that are linearly arranged along a one-dimensional segment, but which also present some degree of bilateral symmetry. This is, in a first approximation, the geometric disposition of our bodily parts along the somatosensory cortex. In a second approximation, we would need to note that surfaces are twodimensional. Exploring scenarios with higher dimensions is a neat extension of this work, but not our focus at the moment.

In our model, a topographic map responds to inputs, *I*, modeled as arrays of *L* bits (with *L* = 100 in all our simulations). An input signal is generated centered around a position *x*^0^ ∈ [0, *L*] such that bits, *I*_*i*_, with *i* = 0, …, *L* − 1, located at position *x*_*i*_ = *i* closer to *x*^0^ have a higher chance of being 1:

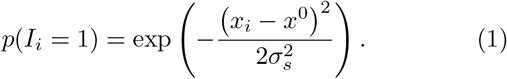

This equation describes an exponential decay with a characteristic scale *σ*_*s*_ (with *σ*_*s*_ = 2 in all our simulations). This means that bits further away from *x*^0^ than *σ*_*s*_ will typically take value 0. This introduces a local spatial correlation between adjacent input bits. The basic topological structure of these input signals is that of a segment of fixed length (Sup. Fig. 1**a-b**). We call such inputs *linearly arranged*. We also refer to them as *fully lateralized* because inputs at one side are different from those at the other.

Additionally, we wish that our inputs also present some level of mirror symmetry, such that bits closer to *L* − *x*^0^ also have a higher chance of being 1. We wish that, if we fold the segment in half like a piece of paper, each input bit’s probability of taking value 1 correlates with its counterpart’s. This is implemented by generating two input signals peaked around *x*^0^, then reversing one of them, and finally transferring each bit with value 1 in the reversed array to the unreversed one with a probability *p*_*T*_ (Sup. Fig. 1**c-d**).

This parameter, *p*_*T*_, controls the level of mirror, or bilateral symmetry in the input. When *p*_*T*_ = 0, no mirror symmetry exists and we recover the fully lateralized case. When *p*_*T*_ = 1, input signals peaked around *x*^0^ are equivalent to those peaked around *L*− *x*^0^, thus obtaining *fully bilateral stimuli*. In [20] it is illustrated how a phase transition seems to exist at some critical value, 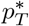, between SOM which, through training, become optimal to represent fully lateralized (for 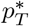) versus fully bilateral (for 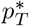) input signals. While bilaterality is important, the most salient topology of the somatosensory map (which we wish to model first) is lateralized. Hence, for our simulations we choose a value 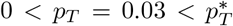 such that trained maps are mostly lateralized but can still display a level of mirror symmetry.

Input signals will be presented sequentially in time to evaluate or train our topographic maps, hence we shall label inputs as *I*(*t*). Responding to these signals are SOM, Kohonen maps [30] that consist of *N* neural units labeled *u*^*j*^, *j* = 1, …, *N* (here, *N* = 200). At each time, each unit is presented with the full input signal and contains a weight, 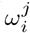, to ponder each of the input bits. The activity of a unit when fed a specific input reads:

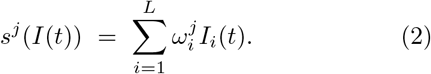

In response to this input, one of the units will have greater activity than the rest. We label this unit *u*^*k*^ and will say that it became “activated”. This active unit hence represents the given stimulus, *I*(*t*). We note this as:

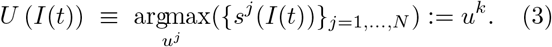

We might present input signals to our SOM just to evaluate which unit is representing stimuli peaked around a given *x*^0^—hence, to find out an activation function *U (x*^0^). Additionally, we might present stimuli to train our network. In that case, after locating the unit that responds to a stimulus, we update all weights according to:

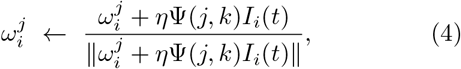

where *η* is a learning rate (here *η* = 0.1) and:

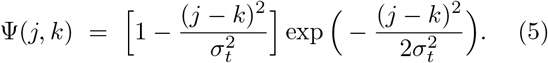

The denominator in Eq. 4 keeps the weights normalized. These two equations mean that the active unit in response to *I*(*t*) and some of its near neighbors become better at representing *I*(*t*), while units outside this neighborhood become worst at responding to *I*(*t*). The neighborhood size is set by Ψ(*j, k*), the second derivative of a Gaußian that constitutes a Mexican hat. The standard deviation of this Gaußian, *σ*_*t*_, determines the size of the neighborhood and represents a local correlation within the neural substrate. In this work, *σ*_*t*_ = 10.

SOM training consisted on the presentation (and consequent weight update) of *T*^*T*^ = 1000 input stimuli thta were independently generated—i.e. each with a randomly, independently drawn *x*^0^. Retraining consisted on the presentation (and consequent weight update) of yet another *T*^*RT*^ = 1000 independently generated input stimuli different from those in the training phase.

Finally, in [20] it was necessary to deal with the emergence of adequate representations. On the one hand, if *σ*_*t*_ and *σ*_*s*_ are very small, representations start forming locally and might not have global consistency. On the other hand, if *σ*_*t*_ and *σ*_*s*_ are large, globally coherent representations emerge, but they are very coarse grained (i.e. they have little spatial resolution in identifying distinctly peaked inputs—because the input themselves present large spatial correlations). Several strategies could be used to tackle this problem. For example, we could anneal the network by reducing *σ*_*t*_ as training proceeds, thus imposing an initial global structure that becomes more refined over time. Some models in the literature dealt with this problem by partly hardwiring input topology into the network structure [36, 40–44].

We chose to *prime* the network by providing, before training, a hint of the global structure of inputs. Therefore, we initialized a mixture of random weights and weights that are optimal to represent fully lateralized inputs. This mixture was weighted by *p*_*P*_, a parameter that controls the priming level. For *p*_*P*_ = 0, fully random weights are chosen; for *p*_*P*_ = 1, SOM start as optimal representations of fully lateralized signals. In [20] a phase transition becomes apparent for some value 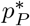 between fragmented and globally coherent representations. In all our experiments, we chose 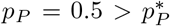, guaranteeing that our SOM are properly formed. Exploring map reorganization for alternative values is an interesting prospect, but in this work we want to focus in a model that captures a realistic description of topographic maps (even if only in its crudest sense).

### B. Modeling stroke injuries

We implement both temporary and permanent focal lesions. *Focal* means that a contiguous set of neural units is affected. This is in opposition to *diffuse* injuries, which would affect non-contiguous units. To determine a focal lesion fully we need to indicate at which position it starts, how many units it affects, and how long the injury persists. We take the superscripts *l* and *λ* (from ‘lesion’, as indexes or injury size) and *S* (from ‘stroke’, to indicate a condition). Thus, the first affected unit is noted *u*^*l*^. A lesion of size *λ* affects units from *u*^*l*^ to *u*^*l*+*λ*−1^. Temporary lesions last for *T*^*S*^ iterations. Trivially, persistent lesions have *T*^*S*^ = ∞.

To simulate permanent lesions, we just ignore units *u*^*l*^ to *u*^*l*+*λ*−1^ from the time of stroke onward. This is achieved in our simulations by clamping their activity, *s*^*j*^, *j* ∈ [*l, l* + *λ* − 1], to a value lower than any activation that can occur naturally in the model. This prevents injured units from ever becoming active. To simulate temporary lesions, we ignore the affected units during *T*^*S*^ iterations after the stroke—again, by clamping their activity to an impossibly low value. We now have an issue regarding training while the injury lasts. When another unit becomes active, the ensuing plasticity also updates the weights of the injured units. We do not know precisely what might happen in real topographic maps. Damaged tissue is probably offline also for learning. Additionally, stroke might incur in damage to the weights that goes beyond the plasticity rules that we implement. For simplicity, we have decided to carry on with the training also for the affected units. This is not an issue in permanent lesions, where clamping to low activity is equivalent to totally removing the affected units (which we also implemented).

We have made experiments changing the location (not shown) and size of our injuries. Regarding location, we did not appreciate important changes in our results. The only deviations that we expected would be in lesions very close to the edges (*l* = 1 or *l* = *N*, but nothing relevant seemed to happen) or around *l* = *N/*2. This second possibility corresponds to the transition from one half of our SOM to the other, with each half corresponding to a simulated hemisphere. Note, however, that in our SOM both hemispheres are contiguous. This makes that good representations often allocate inputs peaked around *x*^0^ = *L/*2 to each hemisphere indistinctly. Lesions around this area would muddle the characterization of contralesional representations, thus we avoided it. In this paper we report lesions starting always at unit *l* = 20 with *λ* ∈ [1, 50]. The largest injuries affect up to unit *u*^69^, thus damage is always confined to the first half of the SOM.

When a good SOM develops, the linear order of input signals is reproduced by the emerging topographic map (Fig. 1**f**). This means that the function *U* (*x*^0^), which tells us which unit represents inputs peaked around *x*^0^, is close to a simple linear function of the kind *U* (*x*^0^) = *bx*^0^, with *b* ≃ 2. When an injury is induced, and some units are damaged, the inputs that were represented by the injured units become *orphan*. Then, they become represented ‘by the next good candidate’. As we will see, this new representation usually lies near the injured area or all the way across the SOM—in the contralesional, mirror symmetric units.

Besides characterizing the lesion within the topographic map (through its location, *u*^*l*^, and size, *λ*) it is also convenient to characterize the *extent* of the lesion in the space of input signals (i.e. within *x*^0^ ∈ [0, *L*]). To do so, we stimulate an injured SOM with input signals peaked at *x*^0^ = 0, then *x*^0^ = 1, *x*^0^ = 2, …, until we locate an input which representation is contralesional. We register it as 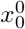. The same process is repeated with signals *x*^0^ = 40, then *x*^0^ = 39, *x*^0^ = 38, …, until, again, we locate a contralesional representation. We register this as 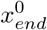 . We call 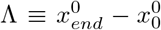 the *extent* of a lesion. The protocol to find the last affected position starts at *x*^0^ = 40 (which in good SOM appears represented around unit *u*^80^; above the last affected unit, *u*^69^) to avoid getting too close to the switch between simulated hemispheres, while making sure that we remain within the first SOM half always.

### C. Rehabilitation protocols

We model some attempts to correct SOM representations. In the literature, we find neurorehabilitation protocols that act directly on neural tissue—e.g. by safely and temporarily inhibiting brain regions using Transcranial Magnetic Stimulation. Other strategies act indirectly on the brain by controlling the kind of input and outputs that a patient has to work with—e.g. increased use of a paretic hand. Our modeled protocols are closer to the later. We alter the frequency with which our SOM encounter input signals across the [0, *L*] segment and look which effect each strategy has on the reorganization of the map.

For each neurorehabilitation experiment, after a simulated stroke we retrain our SOM for *T*^*RT*^ = 1000 iterations as usual. But now, with a 0.5 chance for each iteration, the input is a normal signal (with *x*^0^ randomly drawn from [0, *L*] as usual), or a neurorehabilitating signal drawn from a specific protocol which constraints *x*^0^ to take values within a subset of the input segment. We define Hemisphere (H), Lesion (L), and Border (B) protocols both for *in-lesion* and *out-lesion* conditions as follows:

- **Hemisphere (H)**: Neurorehabilitating signals originate always within a hemisphere, thus *x*^0^ ∈ [0, *L/*2] for the in-lesion condition and *x*^0^ ∈[*L/*2, *L*] for the out-lesion condition.
- **Lesion (L)**: We locate 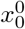and 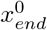 as indicated in the previous section. For the L in-lesion protocol, we draw *x*^0^ from 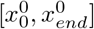. For the L out-lesion protocol we draw *x*^0^ from 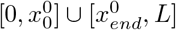.
- **Border (B)**: We locate 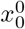 and 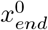 again and use it to generate stimuli that preferentially stimulate the inside or outside border of the lesion. For the B in-lesion protocol we generate stimuli uniformly within 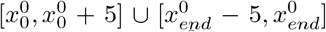. For the B out-lesion protocol the *x*^0^ is uniformly drawn from 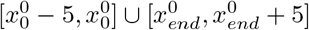.
- **B with offset**: We found that the B out-lesion protocol seemed to interfere with the lesion site too much due to local correlations within input stimuli. We tried similar signals but offset by a margin *m* with respect to the border. These draw *x*^0^ from 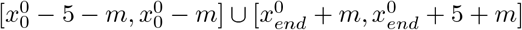.

## III. RESULTS

### A. Single SOM reorganization example

Before unpacking insights from our model, it is worth paying attention to how a single stroked SOM recovers, as shown in Fig. 1**g**. A SOM was trained with *T*^*T*^ = 1000 inputs drawn from an underlying linear arrangement with a hint of mirror symmetry (as specified in Sec. II.A). We simulated a focal lesion starting at *u*^*l*=20^ and spanning *λ* = 45 consecutive units. Retraining with *T*^*RT*^ = 1000 new inputs followed.

We evaluate the resulting topographic map by testing its response, *U (x*^0^), to inputs centered around *x*^0^ ∈ [0, *L*]. The horizontal axis of Fig. 1 shows the activation right after training and before the stroke, *U*^*T*^ *(x*^0^) . The vertical axis shows the activation right after inflicting the stroke, *U*^*S*^ *(x*^0^) (black), and following retraining, *U*^*RT*^ *(x*^0^) (red). The simulated left hemisphere is marked with shadings: blue for the horizontal axis, orange for the vertical one.

The lesion results in a gap in the bottom-left quadrant: since a range of units is missing from the left hemisphere in the damaged SOM, they cannot represent any input. Signals previously represented in that region become *orphan* and must find a representation elsewhere in the SOM. A possibility is that their representation shifts to the right hemisphere—i.e. it becomes contralesional. The upper-left half of the plot contains input signals that, before lesion, were represented in the left hemisphere (blue shading), but after lesion become represented in the right hemisphere (outside orange shading).

In this example we see that most of the orphaned signals become contralesional right after the stroke (black dots in the upper-left quadrant). Some such signals remain contralesional after retraining (red crosses in the same quadrant). But most orphaned inputs are relocated to perilesional units—as shown by red crosses crowding around the lesion area in the bottom-left quadrant.

The upper-right quadrant shows input signals that were consistently represented in the right (unaffected) hemisphere before stroke, immediately after it, and following retraining. This part of the topographic map is virtually unaffected right after the stroke, as marked by the orderly arrangement of black dots—a perfect diagonal would mark *U*^*T*^ (*x*^0^) = *U*^*S*^(*x*^0^) if implemented by black dots and *U*^*T*^ (*x*^0^) = *U*^*RT*^ (*x*^0^) if by red crosses. But after retraining a perturbation arises, as shown by the gap of red crosses in this quadrant. A part of the healthy hemisphere had to make room for orphaned inputs, thus shifting the representation of native, non-orphaned signals. This would be an example of simulated diaschisis, in which a healthy hemisphere’s representation is disrupted by damage to tissue far away.

Our model also shows cases in which all orphaned signals are eventually relocated to the perilesional units in the original hemisphere after retraining. In such cases, disruption to the healthy hemisphere might be avoided (Sup. Fig. 2**a**). But, sometimes, the temporary contralesional representation of orphan stimuli sufficed to have a lasting impact in the healthy hemisphere (Sup. Fig. 2**b**).

### B. A second order phase transition mediates spontaneous SOM reorganization

We ran 1000 simulations with each fixed injury size, *λ* ∈ [1, 50], to find out emerging trends in spontaneous SOM reorganization after stroke. Focusing on *λ* = 45, we compare maps before and right after lesion (Fig. 2**a**), and before lesion and after retraining (Fig. 2**b**). Additionally, Fig. 2**c** shows the changes in units representing each signal, ∆*U*^*S*^(*x*^0^) ≡*U*^*S*^(*x*^0^) *U*^*T*^ − (*x*^0^) (left) and ∆*U*^*RT*^ (*x*^0^) ≡*U*^*RT*^ (*x*^0^) − *U*^*T*^ (*x*^0^) (right) as a function of signal location, *x*^0^.

**FIG. 2.**
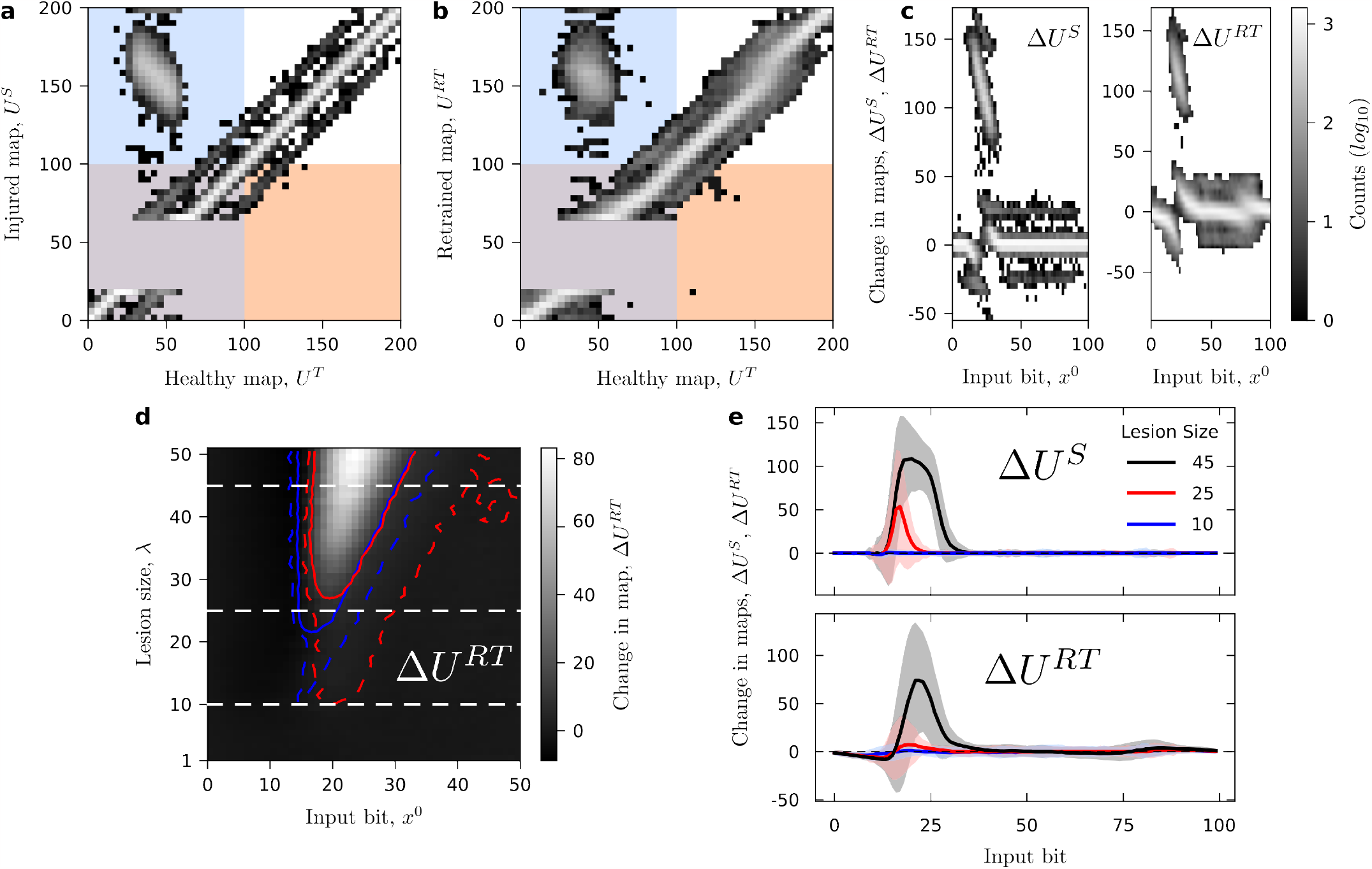
Trends in spontaneous reorganization. Aggregating 1000 reorganizing SOM confirms the patterns that emerge for the individual case. **a** Histogram of shift in representations between healthy maps (horizontal axis) and map right after stroke (vertical axis). Plot is similar to Fig. 1**g**, but now representing the number of simulations within each bin—color bar is at the right side of panel **c**. Shading (blue and orange) similarly indicates hemispheres. **b** Shift in representation between healthy maps and map after stroke and retraining. **c** Shifts in representation ∆*U*^*S*^ ≡*U*^*S*^ − *U*^*T*^ (left) and ∆*U*^*S*^≡ *U*^*RT*^ − *U*^*T*^ (right) as a function of the input’s peak position, *x*^0^. **d** Average shift in representation position, ∆*U*^*RT*^, between healthy maps and maps after stroke and retraining as a function of input’s peak position, *x*^0^, and lesion size, *λ*. Only inputs mapped into the left hemisphere of healthy maps are plotted (see Sup. Fig. 3 for effects on both hemispheres). Contour curves indicate when average representation shift becomes 1 (dashed) and 10 (solid) units. Blue contours are for ∆*U*^*S*^ = 1, 10 (the corresponding histogram is in Sup. Fig. 3**a**) and red contours plot ∆*U*^*RT*^ = 1, 10 (i.e. these are the contour lines of the histogram within this panel). Dashed horizontal white lines indicate cases reported in panel **e. e** Average shift in representation for lesion sizes *λ* = 10 (blue), 25 (red), and 45 (black). Shading indicates standard deviation across simulations.

We identify similar effects as illustrated for a single map: Usually, reorganization involves orphaned inputs and their contralesional counterparts, as well as perilesional units and the signals they represent. Most other input signals remain unaffected, mapped roughly by the same unit, as indicated by the central white diagonal bar in Fig. 2**a-b**, or by the horizontal white bar in Fig. 2**c**. Note, however, the blur around these central features. It extends as far as ∼ 25 units up or down, meaning that there is a likelihood that a few signals outside the affected area nevertheless see their representation displaced by great lengths. This shift is discrete immediately after lesion (as revealed by the gap between the central and shadow bars in Figs. 2**a** and Figs. 2**c** left), and is only partially alleviated by retraining. While rare in our simulations, this would indicate widespread diaschisis.

Back to main features, the extent of contralesional representation of orphaned signals depends on injury size (Fig. 2**d**). We appreciate three qualitatively different regimes: (i) For very small lesions, *λ ≲* 20, all reorganization is perilesional (blue curves in Fig. 2**e**). (ii) Starting at *λ* ∼ 20, and up to moderate injury sizes, *λ ≲*30, representation of affected signals becomes contralesional right after the stroke (red curve in Fig. 2**e**, top); but retraining relocates almost all displaced signals to perilesional areas (Fig. 2**e**, bottom). Finally, for very large lesions, orphan signals become contralesional and, very likely, remain so after retraining (black curves).

Fig. 3**a** shows the fraction, *F*, of simulations displaying contralesional reorganization as a function of lesion size, *λ*. Immediately after stroke (*F*^*S*^, black), the trend reminds strongly of a second order phase transition, displaying an abrupt, yet continuous onset of contralesional reorganization. This further suggests the presence of a critical point. That would mean the existence of a lesion size, *λ*^∗^ ≃ 20, that would poise the topographic map to a state of maximum susceptibility. After the phenomenology of criticality in second order phase transitions, we could also expect fractal patterns of damage, which could obstruct the assessment of impaired tissue. A critical lesion would be quite singular in that slight perturbations could either worsen it terribly, or redirect it with outstanding success. Such heightened susceptibility (captured by responses to slight changes in injury size, *c*_*λ*_ ≡*dF*^*S*^*/dλ*) would be absent well above and well below the critical regime. In such cases, slight nudges would not be as successful, and large interventions would be needed (both to worsen or to restore function).

**FIG. 3.**
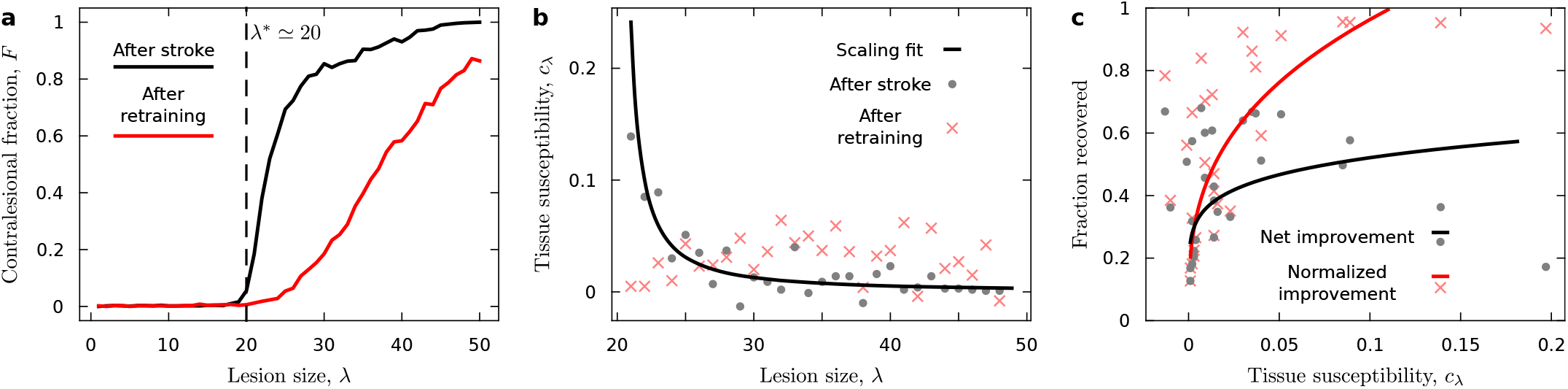
Second order phase transition mediating SOM reorganization after simulated stroke. **a** Fraction of simulations with contralateral representation, *F*, as a function of lesion size, *λ*. The plot right after stroke (*F*^*S*^, black) has the characteristic shape of a second order phase transition: No contralateral representations at all up to a critical value (*λ*^∗^ ≃ 20, dashed vertical black line), followed by a steep increase in *F*^*S*^ at this critical lesion size but with no discontinuity. The phase transition is not present after stroke and retraining (*F*^*RT*^, red). Now the damage increases parsimoniously without any identifiable thresholds separating regimes. **b** The possibility of a second order phase transition with a critical point is supported by studying the susceptibility, *c*_*λ*_ ≡*dF/dλ*. This property measures the SOM’s reaction to infinitesimal changes in lesion size. In second order phase transitions with a critical point, *c*_*λ*_ diverges as a power law as we approach lesions of critical size, *λ* → *λ*^∗^. This is supported by the trend line, *c*_*λ*_∝ (*λ* − *λ*^∗^)^−1.27(*±*0.52)^ (solid black curve). After stroke plus retraining we do not observe this behavior—instead, susceptibility appears rather flat, suggesting a similar response to treatment disregarding of injury size in the chronic phase. **c** Increased susceptibility correlates with better spontaneous reorganization. Gray circles indicate net fraction of recovered cases after retraining, *R* ≡*F*^*S*^ − *F*^*RT*^ . This quantity can be small, e.g. for *λ* = 20 (left-most data point), because for numerical reasons *F*^*S*^ was small in the first place. This improves when we compute the ratio of fraction recovered over originally contralesional representations, *r*≡ *R/F*^*S*^ (light red crosses). Then, for *λ* = 20, we see an almost 100% recovery rate.

The fraction of contralesional representation changes dramatically after retraining (*F*^*RT*^, red in Fig. 2**a**). The phase transition disappears (and thus the critical point), yielding to a more gentle trend. For a range of injury sizes (20 ≲ *λ* ≲ 30), the likelihood that a representation remains contralateral becomes low. This means that, following normal retraining, most orphaned input signals are spontaneously relocated to perilesional units.

The greatest changes appear closer to *λ*^∗^, suggesting that spontaneous relocation might benefit from diverging susceptibilities at criticality (Fig. 3**b**). Fig. 3**c** shows recovery rates, *R* ≡ *F*^*S*^ − *F*^*RT*^ (gray dots) and normalized as *r* ≡ *R/F*^*S*^ (red crosses), as a function of susceptibility, *c*_*λ*_. Trend lines assume power-law scaling (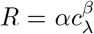 and 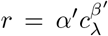), and they show how greater recovery is expected the larger the tissue susceptibility, *c*_*λ*_. For massive lesions (e.g. *λ* ≳ 40), there is a great chance that representation is not relocated ipsilesionally. But, after retraining, this regime is not entered abruptly—no clear thresholds exist.

The picture of this phase transition, and its fading after retraining mediated by large susceptibility, matches the story told by the most recent clinical evidence. On the one hand, a hierarchical model [8, 9] (not a mathematical one, but a qualitative description) has proposed that contralesional relocation strongly depends on lesion size, resulting in regimes similar to those at either side of our phase transition. On the other hand, our model captures the contralateral relocation after stroke seen in the post-acute phase of large injuries, as well as the return to perilesional activity while the contralateral one decreases [11, 23, 24]. We also capture that this recovery is stochastic, with a fraction of cases unable to return to ipsilesional control.

### C. Neurorehabilitation strategies to correct topographic maps

We study now what interventions are most successful in turning contralateral representations into perilesional ones. We focus on a large lesion size, *λ* = 45, for which above 70% of orphan signals remain contralesional after retraining. Some clinical neurorehabilitation strategies include aggressive protocols such as Constraint Induced Movement Therapy (CIMT; which immobilizes a healthy limb to promote usage of the impaired one), or direct inhibition of the healthy hemisphere by Transcranial Magnetic Stimulation (TMS; somehow recreating, in a controlled and harmless way, a lesion in the healthy brain side). These interventions aim at preventing contralesional activity, forcing a reset of ipsilesional tissue. In Sec. IV, we discuss our findings within the context of these protocols.

We test neurorehabilitation strategies that do not interfere directly with the neural tissue (as TMS would do). Instead, we alter the underlying distribution of input signals—i.e. the chance of *x*^0^ across [0, *L*]. Changes in the frequency of tactile inputs can have a profound impact on topographic maps [18, 46]. We wonder whether we can guide brain reorganization by these means alone.

In Sec. II.C we define Hemisphere (H), Lesion (L), and Border (B) protocols both for *in-lesion* and *out-lesion* conditions. These protocols consist in presenting more often stimuli which, in normal circumstances, would have activated the injured hemisphere (H, in-lesion) or its opposite (H, out-lesion), the damaged units (L, in-lesion) or any unit except the damaged ones (L, out-lesion), or the inside (B, in-lesion) or outside (B, out-lesion) border of the damaged region. Additionally, for the B out-lesion protocol, we explore generating input signals with an offset with respect to the lesion borders. We compare these protocols with performance right after stroke (S) and normal retraining (RT).

Fig. 4**a-c** shows, for each of the protocols and the control case (RT), the change in representation location, ∆*U* (*x*^0^), with respect to the SOM before stroke (see Sup. Fig. 4 for variance). We also show the fraction, *F*, of simulations that present any contralesional representation (Fig. 4**d**), and the average extent, Λ, of such representations (Fig. 4**e**) after applying each protocol. Most interventions have a detrimental effect. While some protocols reduce the fraction of simulations that end up with contralesional representations (below the dashed line in Fig. 4**d**), those representations that persist are worsened by all strategies except one (Fig. 4**e**).

**FIG. 4.**
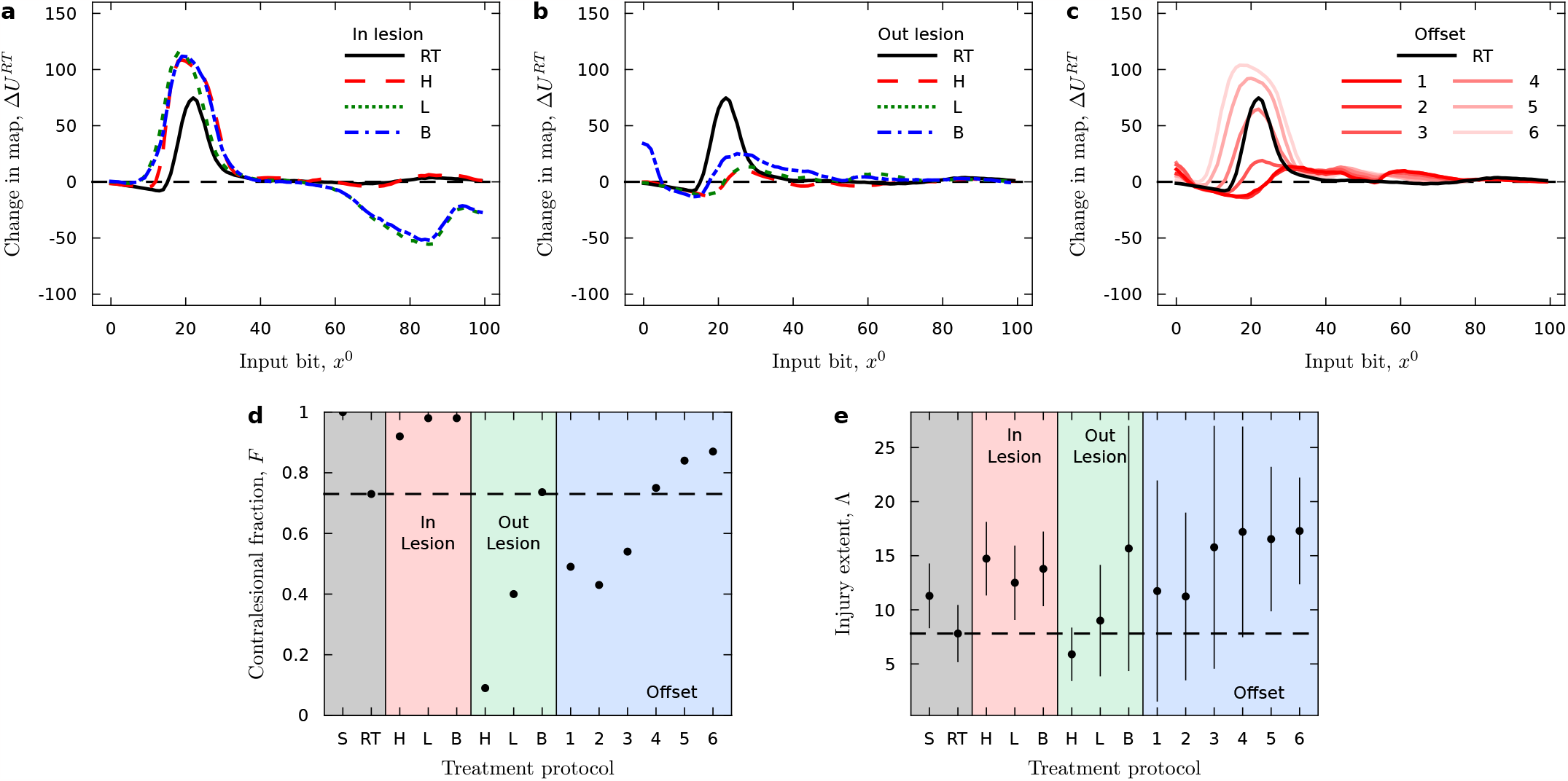
Effect of different neurorehabilitation protocols. **a-c** Shifts in representation between healthy maps and maps after stroke and retraining with a neurorehabilitation protocol. Shifts for normal retraining (without neurorehabilitation) is shown in black for comparison. Same plots with standard deviations are shown in Sup. Fig. 4. **a** Representation shifts for *in-lesion* protocols. **b** Representation shifts for *out-lesion* protocols. **c** Representation shifts for border protocols with an offset. **d** Average fraction of simulations with contralesional representations after each neurorehabilitation protocol and control case (dashed horizontal black line). **e** Extent, Λ, with its standard deviation for persisting contralesional representations after each neurorehabilitation protocol and control case (dashed horizontal black line).

In-lesion protocols L and B also result in great disturbances of healthy representations (Fig. 4**a**). These protocols entail input signals that, in intact SOM, would activate injured units. Once this region is offline, for large lesion sizes, contralesional units become the straightforward option for orphaned signals. Rehabilitation protocols that make immediate use of these inputs cause a prompt reinforcement of their contralesional representations. This hinders perilesional reorganization, and eventually also displaces the native signals of the healthy hemisphere.

The best protocols are, respectively, out-lesion H and L, which explicitly avoid orphan signals. Instead, they stimulate either the contralesional hemisphere (H out-lesion) or any part of the SOM except the injured one (L out-lesion). This last protocol has a chance of stimulating the lesion border, possibly explaining its worse performance. Note, however, that all our protocols are always combined with normal retraining—thus orphan signals are still sporadically presented.

In general, out-lesion protocols that use signals near the lesion border (B and offset strategies) are not great. A few strategies revert a fraction of contralesional representations. But, whenever they fail, they result in much larger lesion extent, Λ. If using inputs too close to the orphan ones, there is a chance, due to local correlations within signals, that they are too similar to some orphan signal. This would entail similar problems as in-lesion protocols. Large offsets seem counterproductive too, as they worsen with respect to regular retraining. An optimal offset happens at *m* = 2, which corresponds to *σ*_*s*_ (i.e. the limit of local correlation within inputs).

### D. Temporary damage suggests delayed timing of ipsilesional reorganization

We now turn to lesions that last a limited time. During such period, damaged units cannot become active. As discussed in section II.B, the plasticity rules remain unchanged—i.e. injured units get their weights updated due to activity elsewhere. We focus on lesions of a fixed size, *λ* = 45, but we study damage that lasts over *T*^*S*^ ∈ [0, 900] iterations. While damaged, our SOM are retrained with usual input stimuli. We wanted to compare performance after a retraining period of *T*^*RT*^ = 1000 steps, but we could consider such time since injury onset or after the injury has subdued. To be thorough, we report performance at a time 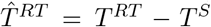 after injury onset (at which point *T*^*RT*^ new retraining inputs have been presented) and at a time *T*^*end*^ = *T*^*RT*^ + *T*^*S*^ (thus marking *T*^*RT*^ iterations after damage subdues); as well as right after the stroke onset, *T*^*S*^.

Fig. 5**a** shows the fraction of contralesional representations as a function of lesion duration, *F* = *F* (*T*^*S*^), for each report time: at *T*^*S*^ (*F*^*S*^, black), at 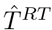 (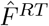, red), and at *T*^*end*^ (*F*^*end*^, blue).

**FIG. 5.**
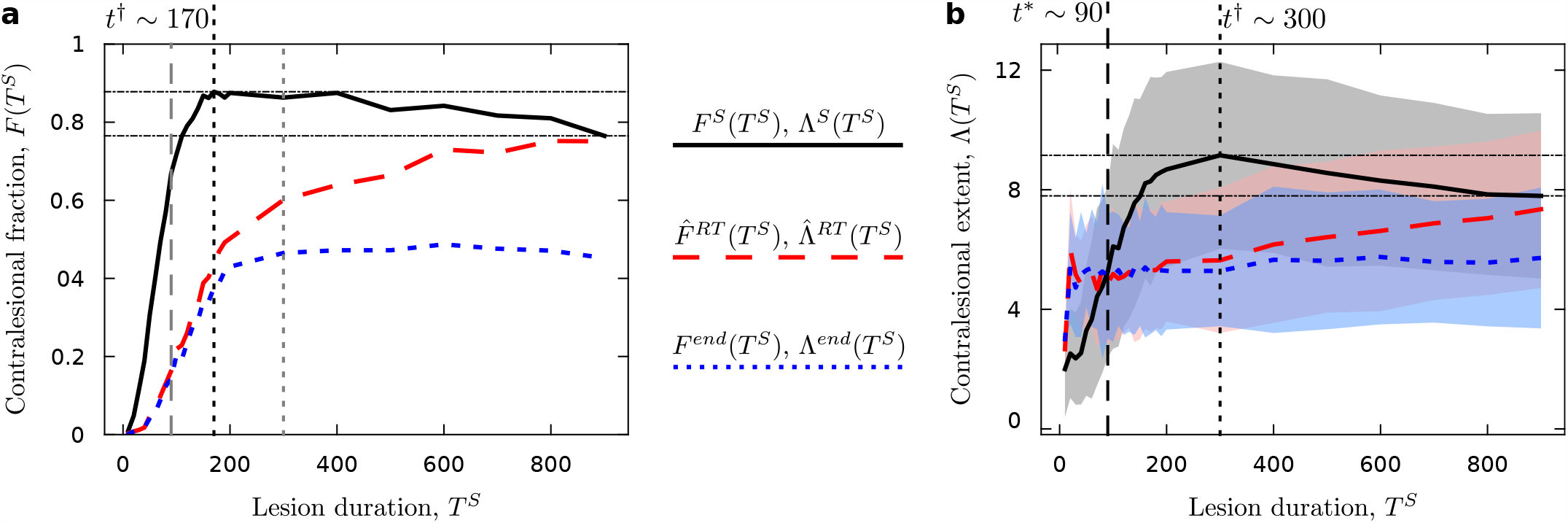
Modeling temporary lesion. **a** Fraction of contralesional representations after temporary lesions of duration *T*^*S*^, as reported right after damage subdues (at *T*^*S*^, solid black), after *T*^*RT*^ = 1000 retraining iterations after stroke onset (at 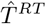, dashed red), and after *T*^*RT*^ retraining iterations after damage subdued (at *T*^*end*^, dotted blue). The global maximum of *F*^*S*^(*T*^*S*^), at *T*^*S*^ = *t*^‡^≡ 170 is marked by a dotted vertical black line, but that curve remains fairly constant within *T*^*S*^ = [*t*^‡^, 400]. The dashed and dotted vertical gray lines correspond to features from panel **b**. Dot-dashed horizontal black lines mark the improvement that longer lesions have over lesions of duration *T*^*S*^ = *t*^‡^. **b** Extent, Λ and standard deviation (shadings) of persistent contralesional representations. The global maximum of Λ^*S*^(*T*^*S*^) happens at *T*^*S*^ = *t*^†^≡ 300 (dotted vertical black line), within the range of maximum *F*^*S*^(*T*^*S*^). Below *T*^*S*^ = *t*^∗^ ≡ 90 (dashed vertical black line), the extent of contralesional representations right after damage subdues is smaller than after retraining.

For *F*^*S*^ we find a rather broad and flat peak. A global maximum appears at *T*^*S*^ = *t*^†^≡ 170 iterations. But, up until *T*^*S*^ = 400 iterations, *F*^*S*^ remains of a rather constant magnitude. For *T*^*S*^ *>* 400 we see a decline. This means that there is a worst-case lesion duration. Any injuries shorter than 170 or longer than 400 iterations result in less contralesional representations right after damage subdues. We speculate that ipsilesional reorganization might be a rather complex process (as it entails the ordered displacement of healthy representations around the lesion site), and that it starts to operate effectively at some point between 170 and 400 iterations after injury onset. This is consistent with the reorganization pattern in permanent lesions (immediate contralesional, then shift to perilesional) found both in our model and in clinical studies. If a lesion lasts longer than the onset of ipsilesional reorganization, it will benefit from this mechanism before the damage subdues, and result in a lower *F* (*T*^*S*^) than at *T*^*S*^ = *t*^†^.

After *T*^*end*^ iterations, there is an improvement with respect to performance at *T*^*S*^ (*F*^*end*^ *< F*^*S*^) and at 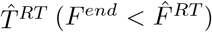. So much so that the effect of the worst-duration vanishes. There are great chances (about 50%) that temporary damage reverts spontaneously with normal activity. *F*^*end*^(*T*^*S*^) reaches a plateau at around *T*^*S*^ = *t*^†^. This suggests that a fraction of simulations reach a seemingly stable configuration with contralesional representations that cannot get undone with regular input signals. Note that, for *T*^*S*^ = 900, total retraining time (during and after damage) equals almost twice usual *T*^*RT*^ ; yet a similar fraction of contralesional cases remains.

The emerging pictures are further supported by Fig. 5**b**, which shows the average and standard deviation of the extent of contralesional reorganization as a function of lesion duration, Λ = Λ(*T*^*S*^), again reported at each relevant time: at *T*^*S*^ (Λ^*S*^, black), at 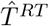 (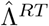, red), and at *T*^*end*^ (Λ^*end*^, blue).

First, Λ^*S*^ has a peak at *T*^*S*^ = *t*^‡^≡ 300 iterations (compatible with the range [170, 400]), again suggesting a delayed onset of spontaneous perilesional reorganization. Second, the existence of stable contralateral configurations is supported by the plateau reached very promptly by Λ^*end*^. This suggests that all sustained contralesional representations after full normal retraining are of a fixed size. Furthermore, Λ^*end*^ *>* Λ^*S*^ for *T*^*S*^ *< t*^∗^ ≡ 90 (in other words, typical contralesional reorganization right after the damage subdues has a smaller extent than after full retraining). It is tempting to say that retraining is harmful in such cases; but, likely, what happens is that only a subset of lesions (those with a persistent contralesional pattern) cannot be reverted by retraining (indeed, Fig. 5**a** shows that a great fraction of cases are recovered by normal retraining for *T*^*S*^ *<* 90).

## IV. DISCUSSION

In this paper we have attempted a minimal mathematical model of brain reorganization after stroke. We build on earlier work on brain hemispherectomy [20] and on the simplest implementation of Self-Organized Maps [30]. Our aim, thus, is to prove that the barest ingredients for large-scale neural plasticity can reproduce dynamics observed in clinical cases; and hence that no additional elements (e.g. circuits overseeing reorganization after damage) are needed.

Our model succeeds in reproducing the most salient, qualitative aspects of spontaneous brain reorganization after a stroke. We find that, for small lesions, reorganization is mainly perilesional; but that, for large enough injuries, contralateral homologues are recruited. This is in accordance with most up-to-date clinical models [8, 9] (which are qualitative; while our description is mathematical, enabling future quantitative research).

For contralesional reorganization in large injuries, we successfully recreate the known dynamics in post-acute and chronic phases observed in the clinic [11, 23, 24]. According to the best accounts available, immediately after a stroke (acute phase) the brain is irresponsive regarding the affected function. This is probably due to unfolding trauma that we do not aim at modeling. In the postacute phase (as soon as 1 to 3 days after stroke), massive damage leads to immediate contralesional activity and strongly reduced ipsilesional activation. This is partly reverted as patients move into a chronic phase (months later): contralesional activity gradually fades out, yielding to reactivation of perilesional tissue. But this is not always the case. In a seemingly stochastic fashion, some stroked brains do not return to ipsilesional control, and affected function remains contralesional in the long term.

Our mathematical model reproduces all these features, including the stochastic retention of contralesional representations in severe cases. We further propose a specific mechanism: the tradeoff between localversus largescale reorganization would be mediated by a second order phase transition with a critical point at a specific injury size, *λ*^∗^. For lesions smaller than *λ*^∗^, reorganization is strictly perilesional—contralateral homologues are never recruited in the first place. For lesions larger than *λ*^∗^, contralesional circuits are the only option immediately available. Simulations with temporary damage suggest that ipsilesional reorganization comes online with a delay with respect to contralateral responses (when these happen, for large lesions). This would explain the shift to contralesional and return to perilesional activity observed in clinical cases and in the model.

Our work is part of an effort to ground neural plasticity in statistical mechanics [20, 47–50]. In classic thermodynamics, we subject a sample of matter (e.g. water) to controlled external conditions termed control parameters (e.g. pressure or temperature). Then we observe how the sample’s configuration changes either gradually (as measured by *susceptibilities*, differential responses to infinitesimal variations of control conditions), or abruptly (as mediated by phase transitions, as when water transforms promptly into ice). If we conceive a neural substrate as a sample of matter, and subject it to relevant controlled conditions (e.g. lesions of specific size), we can bring in the rich mathematical tools and concepts of statistical mechanics to describe how our neural mater gets reorganized.

The phenomenology of critical points and diverging susceptibilities is an example of the relevance of the statistical mechanics framework that our model illustrates. Our simulations show how, immediately after a stroke (when the second order phase transition is present), at the critical lesion size, *λ*^∗^, the neural tissue is optimally susceptible. This means that nudges in the right direction can have a most profound effect to restore a desired configuration. This is quantified by a susceptibility—i.e. the tissue’s response, *c*_*λ*_ ≡ *dF/dλ*, to our control parameter, *λ*. Fig. 3**b** shows *c*_*λ*_ diverging with a power-law scaling, as precisely expected of critical points in second order phase transitions; and Fig. 3**c** shows that increased susceptibility correlates with recovery rates.

These concepts (existence of second order phase transitions, critical points, and susceptibilities) need to be measured by experiments as if brains were thermodynamic samples. Knowledge of the precise location and nature (first or second order) of phase transitions can help us guide brain reorganization as desired. For example, controlled damage of tissue around brain tumors has been used to guide the relocation of neural function, allowing safer tumor resection [51]. Knowing that a second order phase transition might mediate between periand contralesional plasticity would offer strong constraints. Let us speculate: (i) Controlled damage induced to regions smaller than *λ*^∗^ will result in local reorganization. If this is our desired output, we know we can interfere with any area smaller than *λ*^∗^ safely. (ii) If we desire to avoid perilesional plasticity (e.g. because we suspect it affected by the tumor), then interventions larger than *λ*^∗^ would be needed. (iii) At precisely *λ*^∗^, the diverging susceptibility would make neural tissue more easily manageable. Very likely, the timing of the induced damage (which exhaustive modeling falls beyond the scope of this paper) is also of relevance for this speculation.

Our findings are enabled by a tradeoff between perilesional and contralateral reorganization, for which it is indispensable to allow large-scale neural plasticity. Earlier works on focal lesions modeled only local effects [34–39], but we are indebted to a body of work by James Reggia and colleagues, who explored the effects of transcallosal communication in cortical activity [40–44]. Their models dealt with some issues in a slightly different manner than us—e.g., while we encode the spatial arrangement and bilaterality in the input structure, they hard-wire it in the receptive fields of their neural units (which only get inputs from a local neighborhood and from contralateral homologues). They also opt for a 2-D cortical and input structure, and focus in questions slightly different from ours—specifically, in modeling what SOM turn out depending on whether transcallosal connections are inhibitory or excitatory.

Because of this, Reggia et al. explore ranges of parameters that we are not concerned about (e.g. excitatory transcallosal connections, while we stick to the minimalist winner-takes-all dynamics of the original SOM). This produces some results opposed to ours and to clinical evidence—e.g., for excitatory transcallosal feedback, perilesional activity increases immediately after damage and decreases over time [40]. But they also reach findings similar to ours when inhibitory connections are considered— e.g. increased contralesional activity right after damage. Having such features reproduced by different models, both with rather minimal ingredients, strongly suggests that our insights stem from the barest possible dynamics for neural plasticity, and that no additional circuitry is needed for real brains to present the phenomenology summarized from clinical observations.

It is difficult to match our model precisely to the ones by Reggia et al., mostly due to differences in implementation choices and reported measurements. But we tend to think that ours might be similar to some subcase within their wider range of configurations. A more focused study allows us to detect the phase transition mechanism (which Reggia et al. do not find; possibly, we think, because their choice of map and receptive field sizes does not allow them bring such phase transition sharply into focus), as well as the delayed onset of perilesional reorganization that we encounter for temporary lesions. They do not explore neurorehabilitation strategies either—the final point of our discussion.

There is evidence that sustained contralesional activation is harmful to functional recovery after stroke in some cases [11, 24, 28]. But there is also evidence that, in others, contralateral activity is necessary for sustained function [26, 29]. The brain is plastic and complex enough that both possibilities can be accommodated. In any case, understanding the mechanisms that mediate local- versus long-range plasticity (such as our phase transition), and being able to guide brain reorganization as desired is important.

We try different neurorehabilitation protocols and measure their performance as their ability to revert orphan stimuli to ipsilesional representations. Our neurorehabilitation protocols consist in altered probabilities of drawing orphan stimuli. Such stimuli were originally represented by the units damaged by the stroke. Protocols that present orphan stimuli more often (think, in the real world, stimulation of a paretic hand), thus result in a reinforcement of contralesional representations—because they are the representations immediately available after stroke. Protocols that avoid orphan stimuli (and, actually, that over-represent stimuli opposed to the orphan ones) result in an enhanced recovery rate. The reason for this is that over-representation of stimuli opposed to orphan ones tend to expel orphan representation from the healthy brain side. Note, however, that our protocols are combined with normal retraining—hence a small fraction of orphan stimuli is always presented.

This previous finding seems paradoxical. It suggests that to recover a paretic hand we should enhance use of the healthy one—a practice potentially compensatory and maladaptive. There is a very fine point that differentiates our model from real brains. Our model deals exclusively with topographic representations, and in that sense the simulations are clear regarding optimal protocols. But real brains deal with more than topographic maps: On the one hand, a topographic representation links body parts to different cortical areas in the somatosensory and motor cortices. On the other, dedicated circuitry plans and executes movement of each body part—which goes beyond topographic representation. Stroke damage might affect either computational layer, if not both. It is unlikely that injured circuits for executive function are recovered unless specifically exercised, which would demand engagement of orphan activity (e.g., again, attempting to move a paretic hand). But it is also unlikely that these circuits can be properly stimulated if topographic representation miss-directs them to contralesional circuitry. We believe that a complex tradeoff might emerge from these contradictory needs, and that this should be addressed in future research.

It is insightful to check how successful neurorehabilitation strategies have bypassed this issue. Recent research posits CIMT and TMS as promising strategies [52, 53]. The former constrains a healthy limb, for hours on end, to promote only the use of the paretic one. This would lead to a sustained inhibition of the healthy hemisphere, extending beyond the paretic limb and its homologue. An inhibited hemisphere is less likely to attempt control of the paretic hand when exercised—but there is still a chance that it does so, perhaps resulting in a worst performance of the therapy. Two possibilities appear possible: (i) Inhibition is enough to avoid contralesional representations, and the damaged circuits are correctly exercised. (ii) Contralesional representation happens despite CIMT, and healthy circuitry is actually being trained in servicing both hands. This could result in impairment spreading beyond the affected limbs (as it does happen in some stroke cases [22, 53–56]). Neurorehabilitation based on TMS aims at directly shutting off contralesional homologues. This is done in a safe, controlled, and temporary way. In this case, since the relevant contralateral circuitry is specifically shut off, the possibility of topographic misguidance when moving a paretic limb is not possible. We hypothesize, then, that TMS-based protocols are better focused.

To close, besides statistical mechanics, there is another theoretical framework that we would like to highlight. Our main results rely on an underlying conflict between two different topological arrangements or inputs: (i) a simpler, linear arrangement that consists of lateralized signals, and (ii) a certain degree of bilateral symmetry. These and other topological configurations are present across cognitive tasks. They derive from the external world (e.g. topological properties of visual or auditory inputs) or from our evolutionary history (e.g. the bilateral symmetry of the human body and brain, or the linear sorting suggested by contiguity of body parts). Different underlying topologies might be mapped into each other with more or less ease. We think that this is a mathematical problem worth tackling that can result in great insights for neuroscience as well.

## Acknowledgments

The authors wish to thank Susanna Manrubia for her support in carrying out this research. This work grew from enlightening discussions on brain plasticity with Ricard Solé at the Santa Fe Institute (and elsewhere), and with Susanna Manrubia and her extended group of researchers on Complex Systems at the Spanish National Center for Biotechnology (CNB), the Carlos III University, and other institutions in Madrid, Spain.

## Appendix A Supplementary Figures

**SUP FIG. 1.**
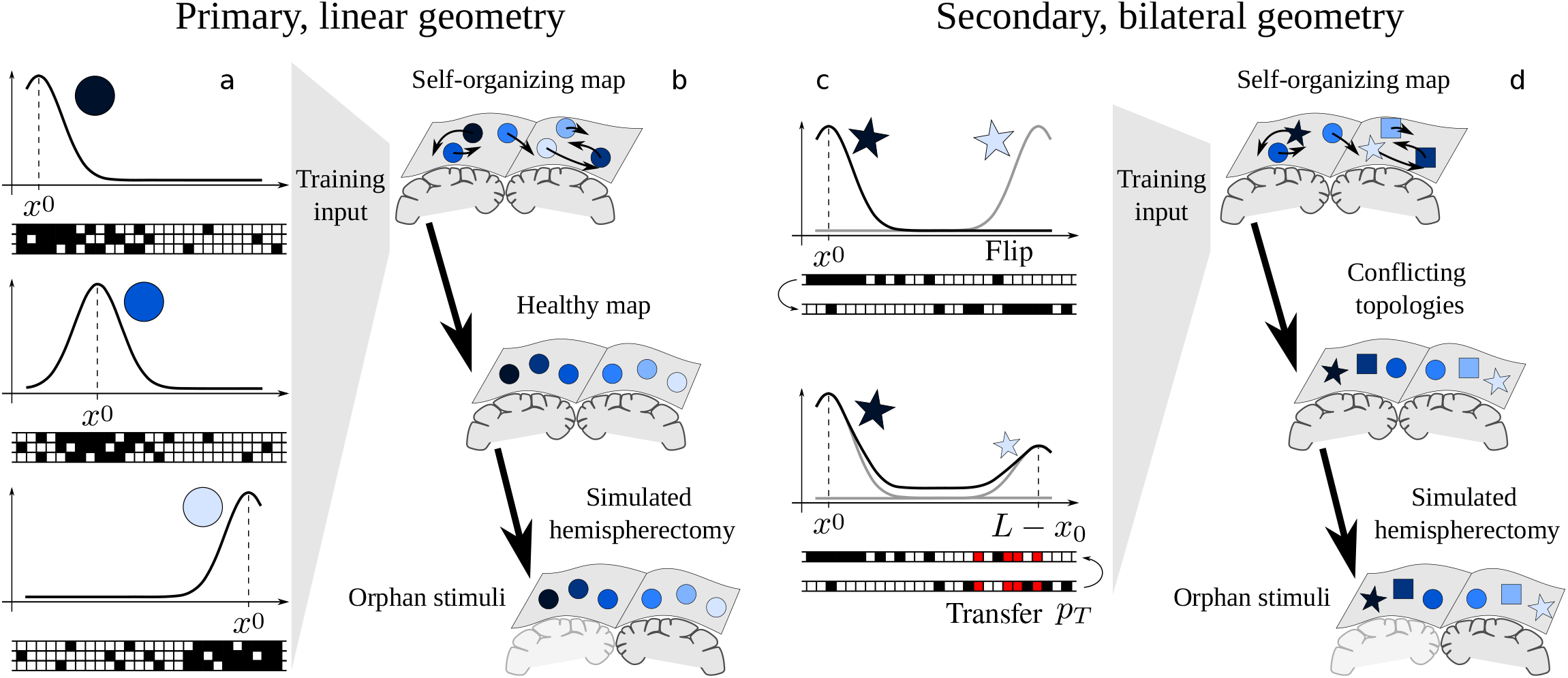
Learning conflicting topological arrangements with self-organized maps. Figure reproduced from [20]. **a** Each input signal consists of a string of bits (shown under the plots). The curves illustrate the probability that a bit takes value 1 (black boxes in the bit strings). This probability peaks at a different position *x*^0^ ∈ [0, *L*] for each signal, thus giving the range of possible inputs an underlying one-dimensional topological disposition. A blue gradient (within circles) further illustrates this contiguity. Three possible signals are shown for each *x*^0^ to highlight their stochasticity. **b** Self-organized maps infer the underlying geometry of the data, thus acquiring an internal geometric structure of their own. **c** We model more complex inputs with the primary linear geometry (gradient-coded) and an additional bilateral symmetry (coded as an interhemispheric correspondence between polygonal shapes). Therefore we create two bit strings as before, flip one of them, and (with a probability *p*_*T*_) transfer some of the bits of the flipped string (red). This results in signals peaked at a position *x*_0_ and at their mirror-symmetric location (*L* − *x*_0_). The bilateral symmetry conflicts with the simple contiguity of linear inputs. **d** Attempts to learn the conflicting geometries might result in SOMs shifting between alternative representations.

**SUP FIG. 2.**
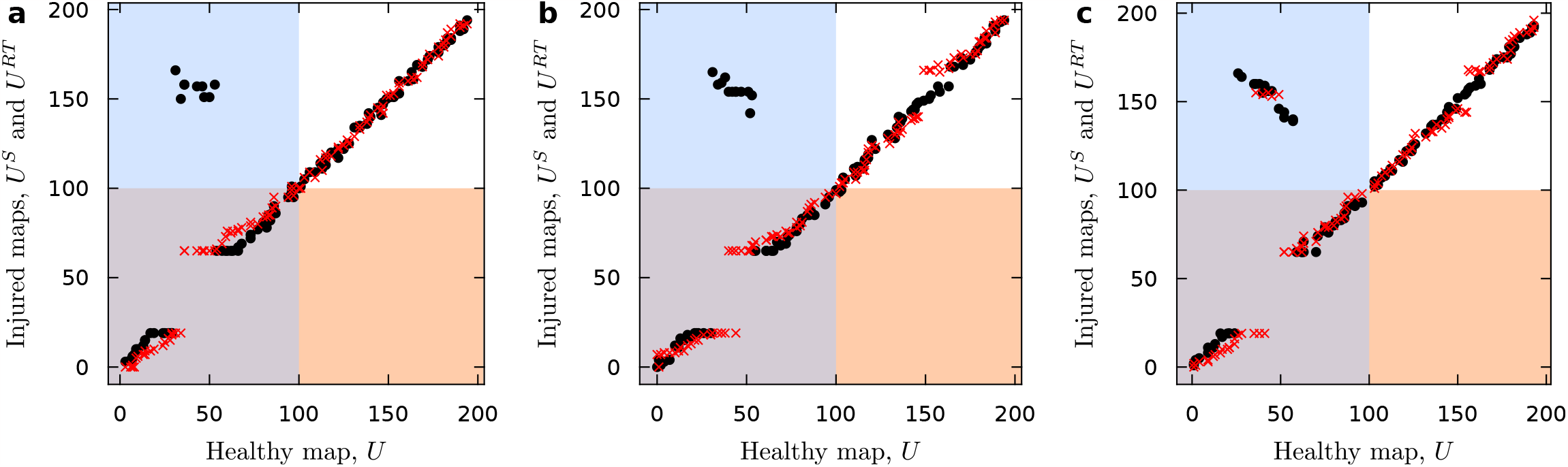
Examples of stroke effects in SOM. **a-b** Cases in which the initial contralesional reorganization is fully corrected into perilesional representations. This can happen without a lasting effect on the contralesional hemisphere (**a**), but the temporary disruption can also result in sustained diaschisis (**b**). **c** Case with persistent contralesional reorganization shown in the main text, reproduced to facilitate comparison.

**SUP FIG. 3.**
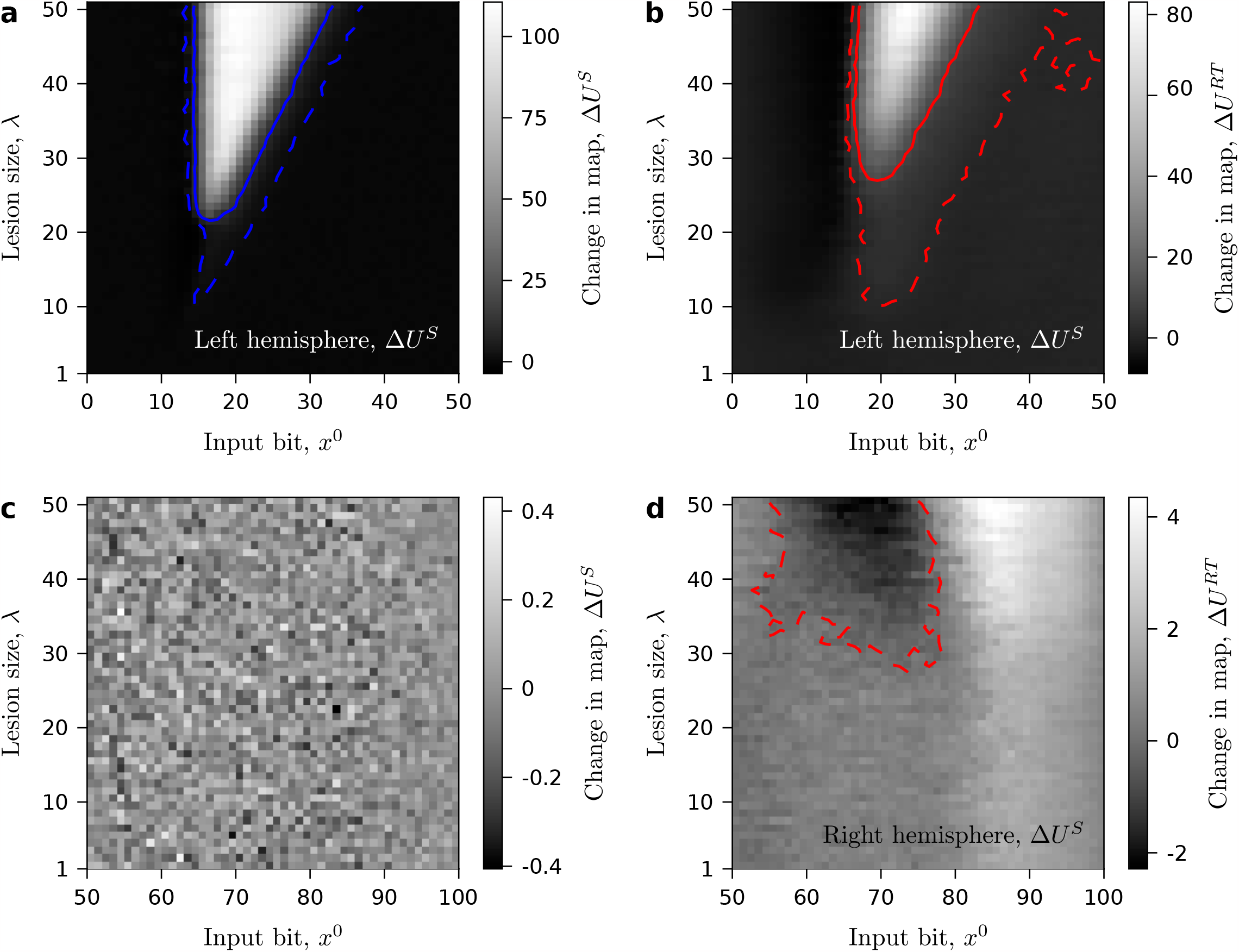
Effects of reorganization in both hemisphere as a function of input’s peak position, *x*^0^, **and lesion size**, *λ*. Dashed contour lines indicate shifts in representations larger than one neural unit, ∆*U >* 1 (i.e. the representation of the corresponding input signal is, in a damaged map, in average, in a different unit than the original in the full healthy map). Solid contour lines indicate shifts in representation larger than 10 units,, ∆*U >* 10. Blue contour lines are used for SOM right after the stroke (**a**) and red contour lines are used for SOM after stroke and retraining (**b** and **d**). **a** Representation shifts in the left (stroked) hemisphere right after damage. Large displacements (well above ∆*U >* 10, likely indicators of contralateral representations) produce a very sharp image. This sharpness in the injury borders is characteristic of threshold-like processes such as phase transitions. **b** Representation shifts in the left hemisphere after stroke and retraining. Large displacements (above ∆*U >* 10) occupy a smaller area now, but persist. Smaller displacements have extended. Perilesional reorganization, including that of signals that had not become orphan, implies that many input signals have been displaced at least one unit in average. **c** Representation shifts in the right hemisphere right after stroke. No pattern emerges. We are observing the stochastic noise of the model, since right after damage the right hemisphere has not been modified (neither by retraining nor by lesion) and signals originally mapped into it remain in place. **d** Representation shifts in the right hemisphere after stroke and retraining. In this case, the effect of diaschisis (perturbation of distal regions due to the original, contralateral lesion) emerges as a pattern of signals displaced from their original representations.

**SUP FIG. 4.**
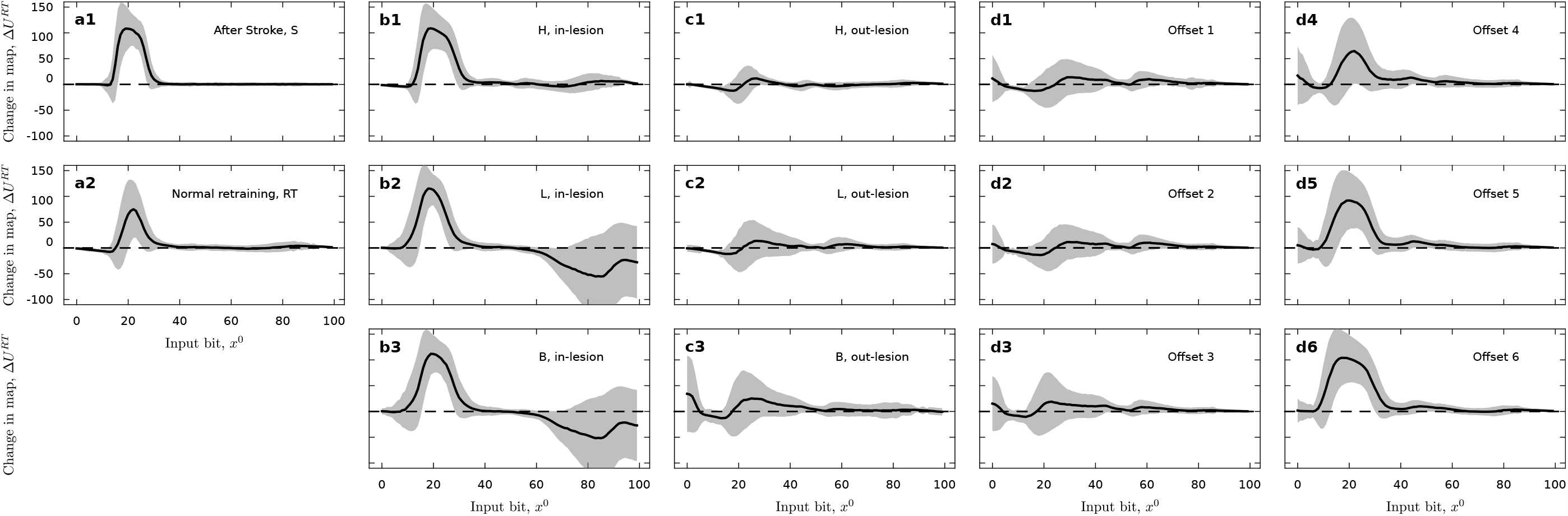
Effect of different neurorehabilitation protocols. Shifts in representation between healthy maps and maps after stroke and retraining with a neurorehabilitation protocol. Shading indicates standard deviations over experiment repetitions. **a** Shifts in representation for control cases right after stroke (**a1**) and after stroke and normal retraining (**a2**). **b** Shifts in representation for in-lesion protocols: hemipshere (**b1**), lesion (**b1**), and border (**b1**). **c** Shifts in representation for out-lesion protocols: hemipshere (**c1**), lesion (**c1**), and border (**c1**). **d** Shifts in representations for border out-lesion protocols with offset.

